# Functional determinants of protein assembly into homomeric complexes

**DOI:** 10.1101/081745

**Authors:** L. Therese Bergendahl, Joseph A. Marsh

## Abstract

Approximately half of proteins with experimentally determined structures can interact with other copies of themselves and assemble into homomeric complexes, the overwhelming majority of which (>96%) are symmetric. Although homomerisation is often assumed to be functionally beneficial and the result of evolutionary selection, there has been little systematic analysis of the relationship between homomer structure and function. Here, utilizing the large numbers of structures and functional annotations now available, we have investigated how proteins that assemble into different types of homomers are associated with different biological functions. We observe that homomers from different symmetry groups are significantly enriched in distinct functions, and can often provide simple physical and geometrical explanations for these associations in regards to substrate recognition or physical environment. One of the strongest associations is the tendency for metabolic enzymes to form dihedral complexes, which we suggest is closely related to allosteric regulation. We provide a physical explanation for why allostery is related to dihedral complexes: it allows for efficient propagation of conformational changes across isologous (*i.e.* symmetric) interfaces. Overall we demonstrate a clear relationship between protein function and homomer symmetry that has important implications for understanding protein evolution, as well as for predicting protein function and quaternary structure.

## Introduction

One of the fundamental challenges in the biological sciences is in understanding the relationship between protein structure and function. This problem is highly relevant not only to the understanding of the evolution of a protein's biological role, but for protein structure and function prediction, protein design, and the prediction of the phenotypic impact of mutations.

Within protein families there is enormous functional diversity, and sequences, folds and domains can all code for different functionalities depending on their surroundings^1–3^. In the dynamic and crowded environment that is a living cell^4,5^, proteins are in constant contact with each other and often carry out their functions as part of larger protein complexes^6–9^. The way that the subunits are organised to form the quaternary structure of a protein complex is a crucial piece of the protein structure-function relationship puzzle, alongside sequences, folds and domains.

Most of the structural information on protein complexes that we have available today is for homomers, *i.e.* protein complexes that are formed by the assembly of multiple copies of a single type of polypeptide chain. Analysis of published X-ray crystal structures shows that roughly 45% of eukaryotic proteins and 60% of prokaryotic proteins can form homomeric complexes^10^. Whilst the high fraction of homomers does reflect biases in protein structure determination, and the fraction of heteromeric complexes (*i.e.* those formed from multiple distinct polypeptide chains) within cells is probably higher, homomerisation is clearly an extremely common biological phenomenon.

Why do proteins assemble into homomeric complexes? Multiple benefits have been proposed^6,11–14^. For example, the construction of a large complex from a number of smaller units, rather than one large peptide chain, provides a means for regulation, as the kinetics of homomer assembly is heavily dependent on monomer concentration. Homomerisation can also provide both coding efficiency and error control, as a smaller genetic space is used compared to a monomeric protein of a similar size. From a purely physical perspective, the formation of a homomeric complex can potentially lead to higher stability, as a larger proportion of protein surface area is buried, minimizing any energetically unfavourable solvent interactions.

Another notable feature of homomeric structures is that the large majority are symmetric. This is striking, considering the irregularity of individual protein molecules and the lack of any underlying symmetry of the individual amino acids. Symmetry is routinely used as a tool to predict both structure and reactivity of many chemical systems^15^ and it has been suggested that, as symmetry contributes to the form of the underlying energy landscape, a symmetric organisation of protein subunits can be associated with increased stability in protein complexes^16,17^. Recent large-scale simulations and directed evolution experiments have concluded that symmetric assemblies are in fact the most energetically stable and favourable, at least for homodimers, and therefore more likely to overcome the entropic cost of complex formation. Hence there is an overrepresentation of symmetric assemblies that are available for evolution^18–20^. The drive towards formation of symmetric structures is also evident when considering ordered protein complex assembly pathways^21–24^. Protein assembly seems to follow an analogous mechanism to protein folding, as the most common homomer assembly intermediates are in their most energetically favourable, often symmetric, conformations^21^.

From the perspective of the specific functions carried out by protein complexes, there are several benefits that could drive the evolution of symmetric homomers. One of the more obvious functional aspects is when active sites are located directly at homomeric interfaces. There are a number of examples of shared active sites, such as that of the homotetrameric dihydropicolinate synthase^25^. As the formation of a protein complex leads to a modification of residue interactions, it also alters the conformational space available, leading to a change in the overall protein dynamics. Access to more conformations can lead to new cooperative mechanisms that facilitate allosteric changes at active sites located at different units in the assembly^26^. The textbook example is haemoglobin, where the oxygen affinity at a specific active site on a monomer in the tetramer is heavily dependent on the coordination of oxygen at the analogous active sites in the complex^27^.

As is the nature of evolutionary questions, there are many difficulties in trying to determine what aspects contribute to homomeric quaternary structures, and functional advantages are often assumed without any direct evidence. Lynch has in fact suggested that the observed distribution of homomer quaternary structures could be explained by purely stochastic, non-adaptive processes^28,29^. A recent coarse-grain modelling study also shows an example of oligomerisation without any obvious functional benefits, emerging purely as a side-effect of the thermodynamic stability of the assemblies^30^. Clearly there is an important open question as to how much of protein homomerisation can be attributed to evolutionary adaptation.

Here we take advantage of the large amount of data that is now available on both protein structure and function, and carry out a systematic analysis on the functional determinants of protein assembly into symmetric homomers. We demonstrate strong relationships between the structural symmetry of complexes and the functions they perform. We also present results suggesting that complex symmetry is linked to specific allosteric functions. These results have important implications for understanding the evolution of protein complexes, as well as potential for being utilised in both the characterisation and prediction of cellular functions in homomeric complexes^28,29^.

## Results and Discussion

### Specific protein functions are enriched in distinct protein symmetry groups

Homomeric protein complexes often assemble into energetically stable structures that are symmetric at the quaternary structure level. This means that group theory can provide a useful metric to characterise and separate most homomeric proteins with respect to a closed or helical symmetry group. The Gene Ontology (GO) project is a large bioinformatics-based initiative with the goal to provide terms for the description of function and characteristics of gene products. The functional annotations of genes in the GO database are based on experiments reported in many different studies, and as such, it is not likely to be uniformly represented across gene products and organisms. However, despite these limitations, investigations of over- and under-representation of GO terms have become one of the main uses of the GO project to date and enrichment analysis are used routinely on sets of genes from various sources with excellent results^31^.

Here we investigate the enrichment of GO terms in a large set of non-redundant protein structures containing only a single type of polypeptide chain (*i.e.* monomers and homomers). Below we first introduce the closed symmetry groups that are commonly seen in homomeric complexes and discuss the top most significantly enriched functions associated with each group. Importantly, we only consider protein structures with at least one GO-term assignment in the Uniprot Gene Ontology Annotation (Uniprot-GOA) Protein Data Bank (PDB) dataset^31^. We determine enrichment of GO terms within the set of annotated structures in order to avoid bias in functionalities that are more highly associated with the PDB than the overall ontology. The enrichments of all GO terms in each symmetry group are provided in Table S1. Because there are many closely related GO terms, we filtered them for redundancy (see Methods) and only discuss the non-redundant GO terms in the main text.

The analyses presented in the main text are based upon a non-redundant set of protein structures generated by clustering structures at the level of 50% sequence identity. To complement this and to show that our results are not due to overrepresentation of certain protein families, we also repeated the analyses at a more strictly filtered set, clustered at the level of domain assignments. These enrichments are also provided in Table S1 and addressed in more detail in the Supplementary Discussion.

### Twofold symmetric homomers

In our analysis, we separate the protein complexes with a single axis of rotational symmetry into those with twofold symmetry (*i.e.* belonging to the C_2_ point group), and cyclic assemblies with higher-order rotational symmetry (C_n, (n>2)_). The logic behind this distinction is that C_2_ dimers contain isologous (*i.e.* symmetric or head-to-head) interfaces, whereas cyclic homomers form ring-like structures via heterologous (*i.e.* asymmetric or head-to-tail) interfaces that allow them to adopt closed ring-like structures (Figure 1a and 1b). Symmetric homodimers comprise the large majority (96.8%) of the C_2_ structures in our dataset, although there are a few higher-order structures (*e.g.* tetramers) with C_2_ symmetry.

**Figure 1:**
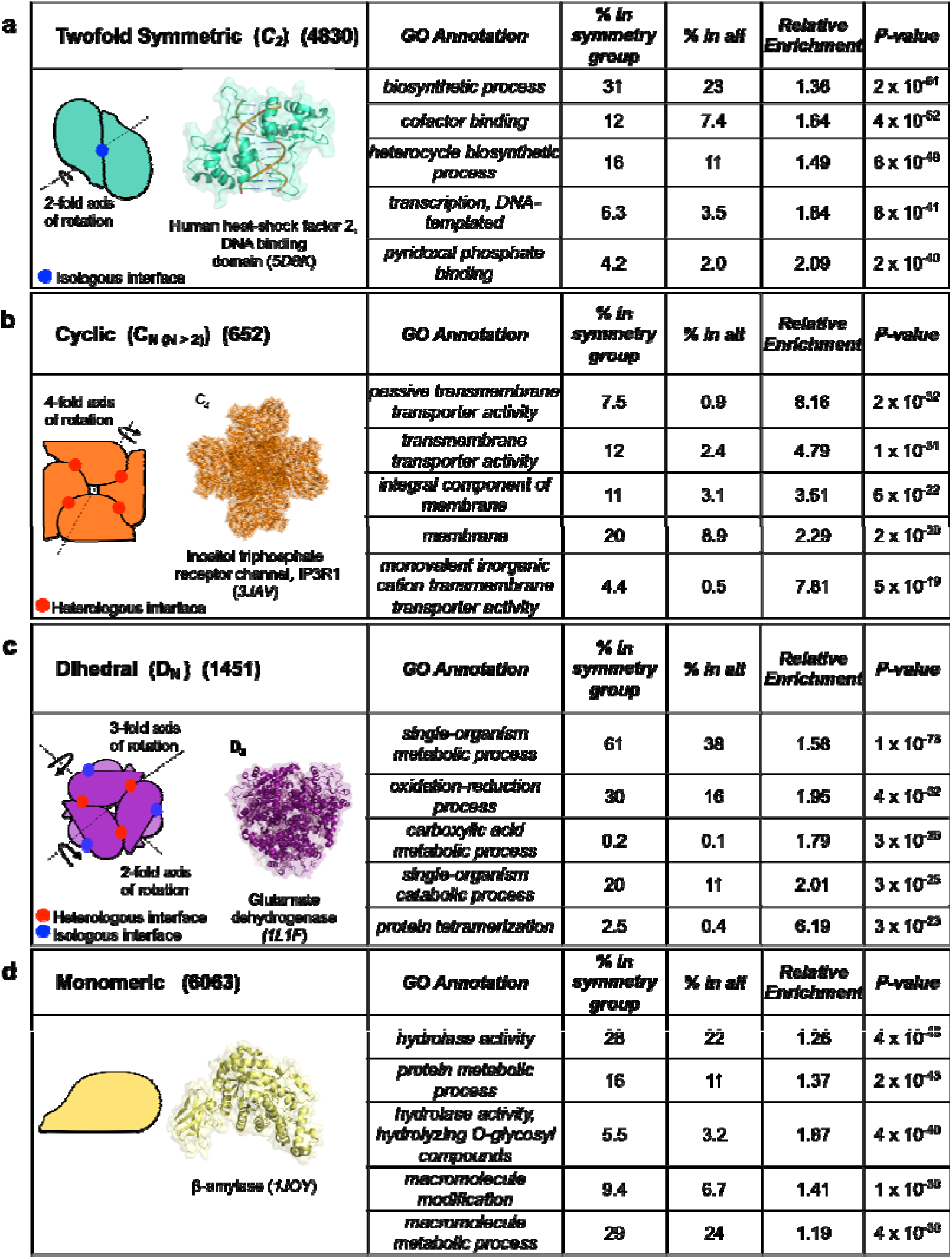
The top five most significant positively enriched non-redundant GO terms within each symmetry group are tabulated with their associated *P*-value from Fisher’s Exact Test. **(a)** Twofold symmetric homomers are associated with annotated functions that involve small and/or twofold symmetric binding partners. Cartoon of a symmetric dimer illustrating the twofold rotation element and isologous interface (blue) with the DNA-binding domain of a heat-shock transcription factor serving as an example. **(b)** Higher-order cyclic protein complexes are required for specialist architectures and are enriched in functional terms involving membrane structures. A C_4_ symmetric complex with the fourfold rotation element and heterologous interfaces (red) highlighted. These interfaces are formed by a head-to-tail orientation of the protein subunits (orange). An inositol-1,4,5-triphosphate activated trans-membrane Ca^2+^ ion channel illustrates an example. **(c)** Protein complexes with dihedral symmetries have a mixture of both isologous and heterologous interfaces and are enriched in metabolic processes. Dihedral complexes in group D_N_ constitutes either *N* symmetric dimers or two symmetric *N*-mers. A D_3_ dehydrogenase illustrates an example of a dimer-of-trimers with heterologous (red) interfaces in the head-to-tail trimers that form a dimer with isologous (blue) interfaces. **(d)** Monomers preferentially act together with large substrates. A β-amylase monomer from B. cereus illustrates an example (yellow).

Our dataset has 4830 non-redundant C_2_ structures, making it by far the most commonly observed symmetry group, comprising 66.3% of homomers. This mirrors previous studies, where small complexes with even numbers of subunits have been found to be much more common in the PDB than larger complexes with odd numbers of subunits^6,32^. An early hypothesis by Monod et al. suggested that an isologous interface is more likely to evolve from a random mutation, as each amino acid occurs twice^33^. It has however since been shown that this effect is somewhat counteracted by the correspondingly low mutation rate of these interface residues, and the prevalence of symmetric dimers seen today is also suggested to be a result of their increased stability^18^.

The C_2_ structures in our set are generally associated with GO functional terms that involve the binding to small substrates. The most significantly enriched term is “*biosynthetic process*” (1.35-fold enrichment in C_2_ structures compared to all structures in our dataset; *P* = 4 x 10^−48^, Fisher’s exact test). This term is defined as the energy demanding conversion of small molecules to more complex substances. One example of the latter is the heterocycle biosynthetic process, which is also significantly associated with the C_2_ symmetric structures in our set (1.49-fold enrichment; *P* = 4 × 10^−40^).

Interestingly, the twofold symmetric structures are also significantly associated with “*DNA-templated transcription*” (1.06-fold enrichment; *P* = 1 × 10^−30^). Although the DNA macromolecule clearly does not qualify as a “small” substrate, many transcription factors, like nuclear receptors, bind to double-stranded DNA at palindromic sequences or inverted repeats^34^, which have local twofold symmetry. Thus, a twofold symmetric homomer provides an ideal mechanism to recognise such a site. This is consistent with, for example, the modular view of transcription factors, where the specific DNA-binding regions can only bind to DNA once a transcription factor dimer has been formed^35^. In Figure 1a, the C_2_ symmetrical DNA-binding domain of human HSF2 is used as an illustration^36^.

### Higher-order cyclic homomers

Our dataset contains 652 higher-order cyclic (C_n(n>2)_) assemblies, comprising 8.9% of homomers. All of the most significantly enriched functionalities in cyclic complexes presented in Figure 1b are related to membranes, *e.g.* “*passive transmembrane transporter activity*” (8.16-fold enrichment; *P* = 2 × 10^−32^). This is unsurprising, as the tendency for membrane proteins to adopt cyclic symmetries has previously been noted^13^. As the cell membrane is effectively two-dimensional, it makes structural sense to find that the cyclic homomers are enriched in this group, forming a defined channel or pore through the membrane with the symmetry axis in its centre. An axis of rotation perpendicular to the membrane plane would imply that the different subunits would be in the same relative position with respect to the membrane bilayer. This set-up has been suggested to be a simplification of the insertion process of the protein in the membrane^13^, although membrane complexes have also been observed to form co-translationally^37^. Whilst the much larger set of C_2_ complexes is found to associate with a large variety of functions, processes and structures, higher-order cyclic complexes are significantly specialised towards functions that require some type of directionality and/or the formation of a channel structure. The recently reported structure of the IP_3_R1 ion channel^38^ is used to illustrate an example in Figure 1b.

### Dihedral homomers

If a protein complex has assembled so that it has two orthogonal axes of rotational symmetry, one being twofold, it is dihedral (Figure 1c) and denoted D_n_, where *n* is the highest order of rotation. These complexes can be thought of as resulting from the assembly of either two C_n_ homomers or *n* C_2_ dimers. In our data, 1451 complexes belong to this point group, and it is therefore the second most common symmetry group, comprising 19.9% of homomers. The twofold symmetry axis means that dihedral complexes always have isologous interfaces but, depending on the mode of assembly, heterologous interfaces can also be present. For example, a “dimer of trimers” structure with D_3_ symmetry will have both isologous and heterologous interfaces, as illustrated with the UDP- glucose dehydrogenase in Figure 1c.

Apart from the functional term “*protein tetramerization*” associated with the formation of the protein complex itself (6.19-fold enrichment; *P* = 3 × 10^−23^), the most significantly enriched GO terms are all related to metabolic processes. For example, more than 60% of the dihedral complexes are associated with the GO term for “*single-organism metabolic process*” (1.58-fold enrichment; *P* = 1 × 10^−73^).

Why might metabolic enzymes have such a strong tendency to form dihedral complexes? One possible reason is that there are advantages to co-localizing multiple different enzymes within a single complex, and dihedral symmetry is a relatively easy way to form a homomers with four or more subunits. For example, a greater number of subunits within a complex can provide a higher concentration of active sites. This can be directly beneficial to the kinetics of its enzymatic function. If the substrate concentration is high, the rate of enzyme activity is a function of not only its rate-constant and enzyme concentration, but also the number of active sites present^39^. The bringing together of many subunits also leads to a generally large enzyme complex being formed from relatively small units. A large protein complex is more likely to be able to provide the correct functional groups for catalysis, have the correct complementarity to its substrate, and also be bulky enough to provide a low dielectric environment for its catalytic process if needed^40^. While large complexes with cyclic or cubic symmetries are possible, it is likely that a dihedral homomers with a given number of subunits is easier to form from an evolutionary perspective^21,23^.

One crucial aspect of enzymatic activity and dihedral complexes that must be mentioned is that it is often beneficial for the enzyme to be able to undergo conformational rearrangements that not only facilitate the process itself, but also change the affinity of the substrates if needed. Bringing together a large number of subunits and forming active sites at interfaces could facilitate this type of allosteric control mechanisms^41,42^. In order to address this important issue, we also investigated our dataset with respect to allostery, as discussed later in the manuscript.

### Cubic homomers

Introducing higher-order rotational symmetry elements leads to so-called cubic complexes. The cubic symmetry groups all have one threefold rotational axis combined with a non-perpendicular axis that can be twofold, as in tetrahedral (T) symmetry, fourfold, for octahedral (O) symmetry, or fivefold, for icosahedral (I) symmetry. These large complexes are assembled from 12, 24 and 60 subunits respectively. These symmetry groups make up only 1.1% of the homomers in our dataset, including 56 tetrahedral, 22 octahedral and 4 icosahedral complexes (most known icosahedral structures are viral capsids, which were excluded from our analysis). The top results from our enriched functions in the cubic set are all associated with homeostasis and metal ion binding in general. This is because, despite our filtering for sequence redundancy, the set contains several octahedral ferritins from evolutionarily diverse organisms. However, even when all but one of the ferritin structures are removed in our dataset filtered for redundancy at the domain level, “*cellular iron ion homeostasis*” is still the most significantly enriched functional term (Table S1 and Supplementary Discussion). This highlights a potential functional advantage of cubic homomers: the formation of large hollow shells ideally suited for storage purposes^6,43,44^. The 24 monomers in ferritin are organised to form a large octahedral capsule capable of storing and transporting iron oxide. The ferritin complex from *Chlorobium tepidum* is shown as an example in Supplementary Figure 1^45^.

### Helical and asymmetric homomers

In our non-redundant dataset of homomers with functional associations, the vast majority (96.2%) belong to a closed symmetry group. Of the exceptions, a tiny fraction (15 or 0.2%) have open helical (H) symmetry, where the symmetry elements contain a rotational axis as well as a translational element along the direction of the rotational axis. These are often found in proteins involved in the formation of long fibres, such as microtubules and actin filaments^46^. The rest (260 or 3.6%) are asymmetric (C_1_) and contain no rotational axes, nor helical symmetry elements.

Both helical and asymmetric complexes are *non-bijective* according to the nomenclature of the recently published “Periodic Table of Protein Complexes”, as sequence-identical subunits are required to exist in topologically non-equivalent positions^23^. Interestingly, however, the majority of these non-bijective homomers are the result of quaternary structure assignment errors^23,47^, often due to the PDB biological assembly erroneously being defined to be the same as the asymmetric unit. Thus, we must be cautious interpreting enrichments in this group, as most quaternary structures are likely to be erroneous. Despite this, it is interesting to note the most significantly enriched functional term for these complexes is “*signal transducer activity*” (3.44-fold enrichment, *P* = 5 × 10^−6^; Figure S1), as the prominent role of asymmetry in signalling processes has previously been noted^48,49^. The second most significantly enriched term is “*DNA binding*”(2.09-fold enrichment, *P* = 1 × 10^−5^). This can be rationalised by the 2:1 stoichiometry of the interaction between a dimeric DNA-binding protein (*e.g.* transcription factor) and the double-stranded DNA, which will be inherently asymmetric unless there is twofold symmetry and the DNA level^10^.

### Monomers

Finally, we consider the functions associated with monomers, *i.e.* structures made up of single peptides that do not assemble into homomers (Figure 1d). In our non-redundant set, 6063 proteins were monomeric. Thus, in our full dataset of protein structures containing only a single type of polypeptide chain, 54.6% are homomeric and 45.4% are monomeric, supporting the idea that most proteins can form homomers – at least of those that can be crystallised alone.

The most significantly enriched functions in monomers are “*hydrolase activity*” (1.26-fold enrichment, *P* = 4 × 10^−48^) and “*protein metabolic process*” (1.37-fold enrichment, *P* = 2 × 10^−43^). We investigated this further by calculating the enrichment of monomeric structures within each hydrolase subclass, according to the Enzyme Commission classification scheme^50^ (Figure S2). Monomers are negatively associated with hydrolase activity on amide bonds that are not associated to peptides (*P* = 4.51 × 10^−5^) and positively enriched for glycosylase activity (*P* = 1.58 × 10^−4^). Overall, this suggests a preferential binding to large hydrolase substrates such as peptides and oligosaccharides.

Another strongly enriched term in monomers is “*macromolecule modification*” (1.41-fold enrichment, *P* = 1 × 10^−30^), which refers to modification of a biological macromolecule, such as a polypeptide or polysaccharide, which alters its properties. Thus, the terms most significantly associated with monomers are all clearly related to processes, components and functionalities that involve large macromolecular substrates. This makes logical sense from a structural point of view, especially comparing with complexes of higher-order symmetry, as a monomer would be more likely to be able to accommodate large substrates. The only functionality that involved binding to a large molecule that was enriched in the twofold symmetric homomers was “*DNA binding*”, which is related to the dimeric and often symmetric nature of DNA. In contrast, if we look through our full enrichment analysis for “*RNA binding”*, we do indeed find it ranked #13 of the positively enriched functions in the monomeric structures (1.42-fold enrichment, *P* = 5 × 10^−16^).

Interestingly, the most significantly negatively enriched functions in the monomer set were related to metabolic processes involving small molecules and organic substances (Table S1). This reflects the fact that these substrates tend to be processed by homomeric enzymes, as shown earlier.

Finally, because monomers comprise such a large fraction of our dataset, we also repeated all enrichment analyses for the different homomers symmetry groups with a dataset that excluded monomer structures (Table S1). In general, the same enrichments were observed both considering and excluding the monomers (see Supplementary Discussion).

### Influence of interface size on functional associations

In the above analysis, homomers have been grouped by symmetry, with no further consideration for structural properties. However, we know that the size of intersubunit interfaces can also be related to function: larger interfaces tend to be stronger and are more likely to be obligate, whereas smaller interfaces tend to be weaker and are more likely to be transient^51–53^. Therefore, we also performed an analysis where we split the homomers from each symmetry group into two equally sized sets: one containing complexes with larger interfaces and one with smaller interfaces. We then compared the relative enrichment of each functional term between the two sets (Table S1).

In general, the functional terms we have identified as being associated with each symmetry group above tend to be enriched in complexes with larger interfaces. This probably reflects the fact that structures with larger interfaces are more likely to represent the biologically relevant quaternary structure within the cell, whereas complexes with smaller interfaces are more likely to be formed only transiently within the cell, or be artefacts of crystallisation.

There are a few functional terms clearly enriched in complexes with smaller interfaces. For example, “*hydrolase activity*” is enriched in C_2_ homomers with smaller interfaces. Since we observed above that hydrolases are overrepresented in monomers, this probably indicates that C_2_ homomers with small interfaces are more likely to be monomeric within the cell. In addition, “*signal transduction*” is also enriched in C_2_ homomers with smaller interfaces, probably reflecting the prevalence of transient dimerization in signalling pathways^11^.

### Allostery is related to the types of intersubunit interfaces formed

The term allostery is generally used to describe a dynamic, cooperative process that involves a change at one site of a protein as a result of an event, such as binding of a substrate, at a distant site of the protein. Allostery has often been associated with symmetric complexes and symmetry is integral to the first and most widely used formulation of allostery, the Monod-Wyman-Changeaux (MWC) model^33^. In this model, allosteric movements are explained as taking place in a concerted manner where the allowed conformations in the protein are restricted to those where the subunits that are allosterically coupled remains symmetric with respect to each other. Despite its simplicity, the MWC model has been remarkably successful at explaining experimentally observed allostery in proteins^54^. Although allostery does occur in many monomeric proteins^26^, symmetry provides a very simple mechanism for allosteric regulation that has been commonly utilised throughout evolution. Therefore, we were interested in investigating how allostery is associated with different types of protein symmetry.

First, we need a way to identify putative allosteric proteins. Although it has been suggested that all dynamic proteins are allosteric to an extent^55^, here we are interested in proteins that have been actually noted as being allosteric, typically due to the biochemical observation of cooperative regulatory behaviour. To do this, we divided our set of structures into those that are putatively allosteric and those that have not been annotated as being allosteric. Although there are no GO annotations for allostery, we used the protein complexes included in the Allosteric Database^56^ as well as a keyword search to identify further putative allosteric proteins (see Methods). While this approach is not perfect, it does allow us to define a set of proteins that should be highly enriched in allostery. We then assessed whether allostery is associated with different symmetry groups, as presented in Figure 2a. Interestingly, we find that dihedral homomers are by far the most strongly enriched in allostery, with a slight enrichment also observed for the C_2_ homomers. Importantly this overall enrichment profile of allostery is still evident when we control for metabolic enzymes, as illustrated in Figure 2b, suggesting that this trend is not simply due to the fact that metabolic enzymes tend to be allosterically regulated.

**Figure 2:**
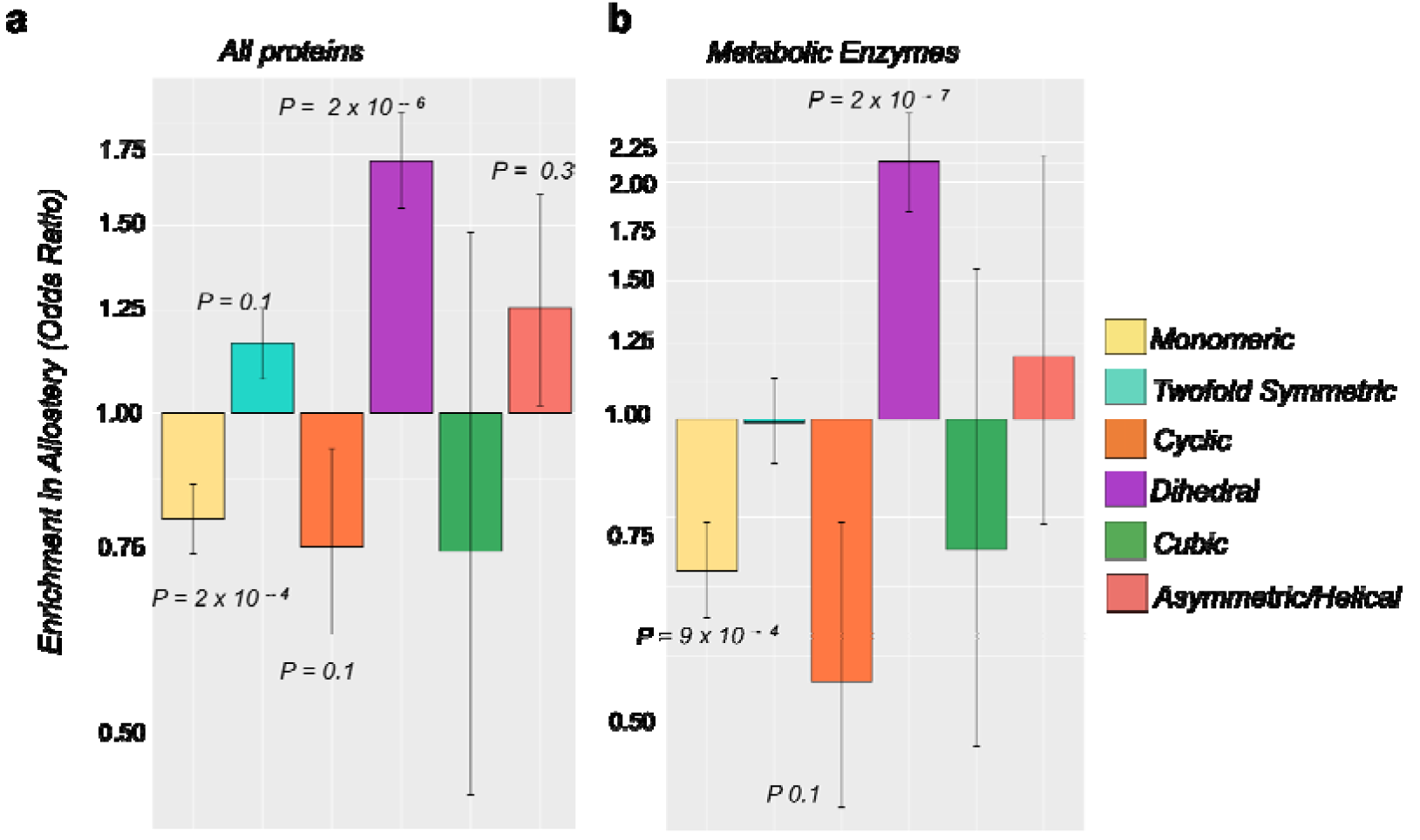
Enrichment of allostery in homomers from different symmetry groups. **(a)** Enrichment by complexes with closed symmetry in the subset of allosteric proteins**. (b)** A similar enrichment profile is seen when controlling for metabolic enzymes. The enrichment is calculated as the difference between the fraction of complexes in the specific set compared to the whole set of homomers. The enrichment is presented as the odds ratio, plotted on a logarithmic axis. *P*-values are calculated with Fisher’s exact test and error bars represent 68% melded binomial confidence intervals.

In principle, any symmetric arrangement of subunits should be compatible with allostery, so there is a question as to why it is enriched in dihedral, and to a lesser extent C_2_, complexes. Interestingly, these two symmetry groups are both associated with isologous protein interfaces, whereas cyclic complexes have only heterologous interfaces. Therefore, to further investigate whether there is a relationship between isologous interfaces and allostery, we separated the dihedral homomers into tetramers (D_2_), which have exclusively isologous interfaces, and those with six or more subunits (D_*n*(*n*>2)_), which can possess a mixture of both isologous and heterologous interfaces, as illustrated in Figure 3a. Analysing the enrichment in allostery in these two sets of dihedral homomers shows a significant enrichment in allostery for both groups compared to the rest of the homomers, but the enrichment is much stronger for the D_2_ set (Figure 3b). This suggests that isologous interfaces drive the enrichment of allostery in dihedral complexes.

**Figure 3:**
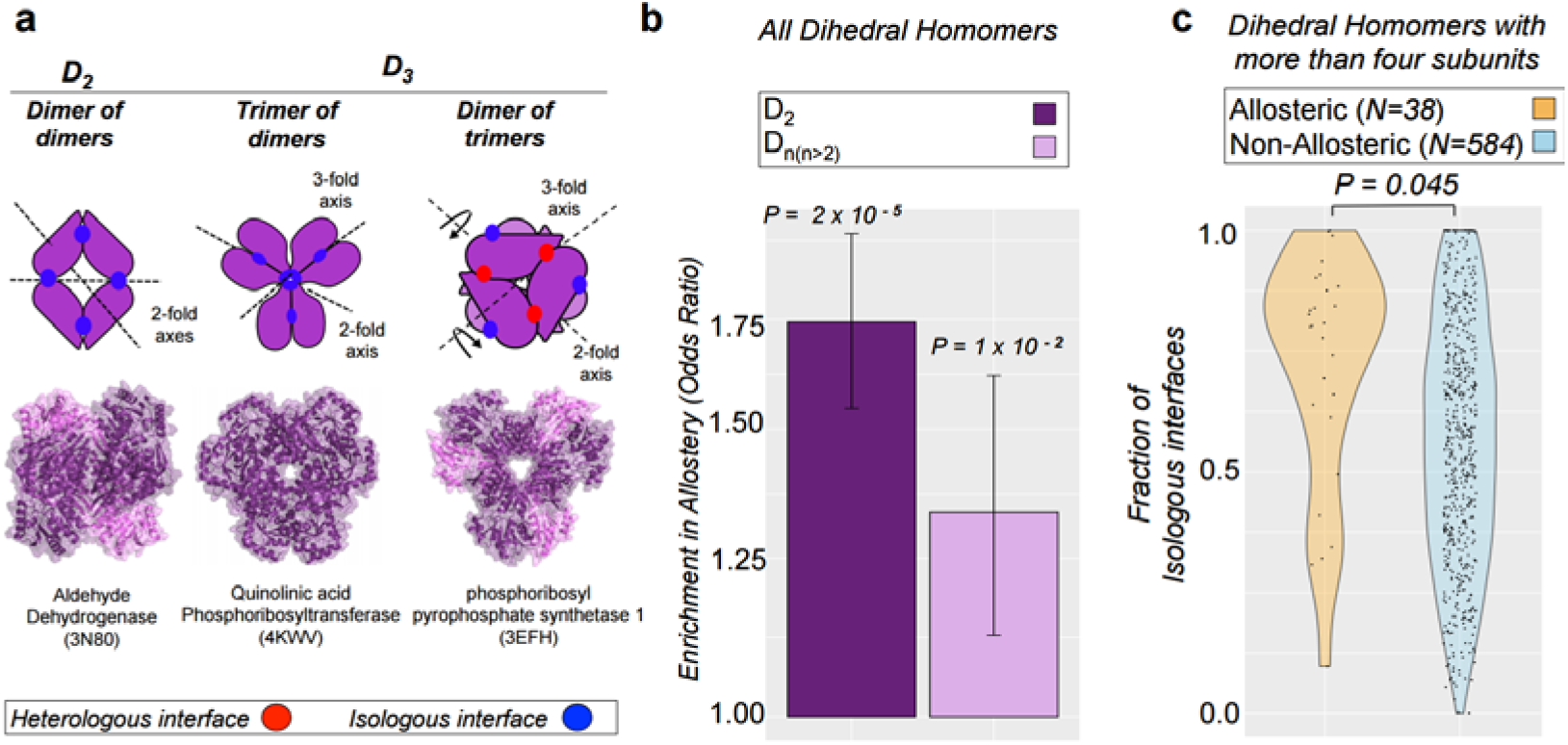
Allostery is associated with the present of isologous intersubunit interfaces. **(a)** Cartoon representation of the intersubunit interactions D_2_ and D_3_ complexes. D_2_ complexes have two perpendicular twofold symmetry axes and therefore have only isologous interfaces, whereas D_3_ complexes can have either all isologous interfaces or a mixture of isologous and heterologous interfaces. **(b)** Comparison of the enrichment in allostery between dihedral tetramers (D_2_) vs. dihedral homomers with six or more subunits (D_n(n>2)_). *P*-values are calculated with Fisher’s exact test and error bars represent 68% melded binomial confidence intervals. **(c)** Density plot illustrating the distribution of isologous interfaces in the allosteric (orange) and non-allosteric (light blue) sets of dihedral complexes with more than four subunits. The *P*-value is from the two-sample Wilcoxon test.

We can also look specifically at the dihedral complexes with six or more subunits (D_*n*(*n*>2)_). Since they can have a mixture of isologous and heterologous interfaces, we wondered whether allosteric complexes would tend to contain a greater proportion of isologous interface than non-allosteric complexes. For every D_*n*(*n*>2_) homomer, we calculated the fraction of the total amount of intersubunit interface formed that is isologous and plotted the distributions for allosteric and non-allosteric complexes in Figure 3c. Interestingly, we observe a significant tendency for allosteric complexes to have a greater fraction of isologous interfaces (*P* = 0.045, Wilcoxon rank-sum test), further supporting the idea that isologous interfaces are conducive to allostery.

Why would allostery be associated with isologous interfaces? We hypothesise that this is due to the symmetric nature of isologous, or “head-to-head” interfaces, in which two identical protein surfaces interact with each other. If a conformational perturbation occurs on one side of an isologous interface, specific to one subunit, then the symmetry of the interface will be broken, but can easily be restored by an identical change occurring on the other side of the interface. Thus, an isologous interface provides a simple physical mechanism to propagate conformational changes from one subunit to another. In contrast, for a heterologous or “head-to-tail” interface, in which two different surfaces interact, it is more difficult to induce an identical conformational change from one subunit to another.

The apparent association between allostery and dihedral complexes was previously noted by Goodsell and Olson^6^. They describe two mechanisms by which allostery often occurs: rotation of two rings with respect to each other and pincher-like motions, that are both compatible with dihedral but not cyclic symmetry. In contrast, they suggest that allostery is difficult in cyclic complexes as it must propagate one subunit at a time around the ring. Our results here expand upon this, showing that allostery is facilitated by the nature of isologous interfaces, and that even when we consider only dihedral complexes, those with more isologous interface are more likely to be allosteric.

## Conclusions

By using the large amount of high quality protein structure data that are now available in combination with functional annotation data, we have been able to gain essential insight into the functional specialisations of homomeric protein complexes. We have shown that protein subunits tend to organise themselves into homomeric assemblies with clear functional preference depending on their overall symmetry, which can be mostly explained by simple geometrical arguments. These functional benefits are likely to have contributed towards positive evolutionary selection for protein homomerisation. However, it is important to emphasise that these represent overall trends. While our results clearly support the idea that homomer quaternary structure can be functionally beneficial, they do not exclude the likelihood that differences in quaternary structure may often be non-adaptive^28,29^.

It is also important to point out that, even if a protein does not comply with any symmetry conditions as a whole, its functional module (for example a DNA-binding- or catalytic site) might be related to the homomeric symmetry groups studied here. Any function that is specifically associated to a symmetry group in this study therefore also has the potential to help characterise, for example, asymmetric heteromeric complexes with a symmetric catalytic site and complexes with pseudosymmetry.

Finally, this work also shows that the formation of isologous interfaces between subunits of the protein complex can provide specific functional advantages. Essentially, these types of interfaces provide a structural mechanism for transmitting cooperative conformational changes across multiple subunits of the complex, which can contribute to allosteric mechanisms. These results provide an added degree of information that can have future practical advances when it comes to the prediction of both function and malfunction of protein complexes.

## Methods

### Structural data

Starting with the full set of protein crystal structures in the PDB on 2015-03-19, we selected all of those containing only a single type of polypeptide chain containing at least 30 residues, *i.e.* all of the monomers and homomers. Viral structures, which were dominated by capsids with icosahedral symmetry, and structures with known quaternary structure misassignments^47^ were excluded. For each structure, we used the first biological assembly (*i.e.* pdb1 file). In order to compile a non-redundant set of structures the chains were clustered into groups according to their sequence similarities and selecting one representative of each cluster. Monomers and homers were then clustered at the level of 50% sequence identity, and only a single structure was used from each cluster. This resulted in a set of 13353 non-redundant structures of either monomers or homomers with functional assignments. Symmetry assignments were taken directly from the PDB. For the comparison of complexes with larger *vs* smaller interfaces, complexes were split into two equally sized sets for each symmetry group, bigger than and smaller than the media per-subunit interface size. Interface symmetry was calculated from the dihedral homomer structures as was done previously^23,57^. Briefly, interfaces were classified as isologous if the correlation between the residue-specific buried surface area for each subunit in an interacting pair was >0.7. All complexes are provided in Table S2.

For an even more strictly filtered dataset, we clustered structures using SUPERFAMILY domain assignments^58^ instead of sequence identify. SUPERFAMILY predicted assignments were used rather than SCOP assignments because they are available for many more structures. Analysis of this dataset containing 5431 structures revealed enrichment of similar functionalities, and is presented in Table S1 and the Supplementary Discussion.

### Functional data

Functional assignments were taken from the Uniprot Gene Ontology Annotation (Uniprot-GOA) PDB dataset (2014-09-27 release), which provides GO functional annotations to PDB chains^31^. In this dataset, a GO term is associated to a chain that maps with at least 90% identity to a UniProt knowledgebase entry. Calculating GO enrichment only within the structure annotated GO term set avoids bias against type of functional terms that are enriched in the PDB. This set was then extended by us to include all associated GO terms (terms that include the *is_a* and *part_of* association in the ontology) from the core ontology database, as curated by the Gene Ontology Consortium. Each GO term was then analysed with respect to the non-viral structures in our set, and the relative ratio of enrichment of protein symmetry groups within each functional term of our set (which were those functions associated with structures in the PDB, rather than the full ontology) were evaluated and tested for significance using Fisher’s exact test, where the null hypothesis is that the odds ratio equals one. Because there are many very similar GO terms, we filtered the GO terms for redundancy, removing terms that share an association with >50% of the same proteins from our structural set. All GO terms, including those we classified as redundant, are provided in Table S1. In order to avoid any bias in our enrichment set caused by there being a larger percentage of monomeric structures, the above analysis was also carried out without monomeric proteins. The general trend in enriched functional terms as discussed in this analysis still holds in the set without monomers, as presented in Table S1. Confidence intervals, at 68%, for the enrichments were calculated using a melded binomial procedure, known to provide a good match for *P*-values of a Fisher’s exact test^59^.

### Identification of allosteric proteins

To identify putatively allosteric proteins, we searched PubMed for the terms “allostery”, “allosteric” and “allosterism”, and then identified PDB structures associated with these abstracts through the NCBI Structure site (http://www.ncbi.nlm.nih.gov/structure). We also accessed all the proteins associated with allostery in the Allosteric Database^56^. Allosteric proteins are listed in Table S2.

## Acknowledgements

We thank György Abrusán and Jonathan Wells for helpful comments on the manuscript. J.M. is supported by a Medical Research Council Career Development Award (MR/M02122X/1).

## Additional Information

### Author contribution

T.B. performed the computational analyses and prepared the figures with supervision from J.M. Both T.B. and J.M. wrote the manuscript text.

